# Image-based phenotypic profiling of a chemogenomic screening library identifies novel druggable targets in the EGFR-pathway

**DOI:** 10.1101/2021.04.16.440090

**Authors:** Kenji Tanabe

**Author notes:** Correspondence should be addressed to Kenji Tanabe. Medical Research Institute, Tokyo Women’s Medical University, 8-1, Kawada-cho, Shinjuku-ku, Tokyo 162-8666, Japan.

## Abstract

The gene encoding epidermal growth factor receptor (EGFR) is a major driver gene in cancer. Many drugs targeting EGFR-associated molecules have been developed, yet many have failed in clinical trials due to a lack of efficacy and/or unexpected side effects. In this study, I used image-based phenotypic profiling to screen a pharmacologically active compound library with the aim of identifying new druggable targets in the EGFR pathway. As anticipated, the phenotypic screen identified compounds that produce phenotypes resulting from targeting a known specific molecule or pathway. The assay also showed that compounds with diverse known mechanisms of action produced similar, EGFR-related cellular phenotypes. Biochemical assays revealed that those compounds share a previously unappreciated common target/pathway, showing that the image-based assay can identify new target molecules that are independent of the compound’s known target. Further experiments showed that ROCK1 and PSMD2 are novel druggable targets within the EGFR pathway.

## Introduction

Epidermal growth factor receptor (EGFR) is a prototype receptor for receptor tyrosine kinases. Many studies have investigated EGFR downstream signaling, and have uncovered molecular mechanisms essential for both EGFR membrane trafficking and signal transduction (Bergeron et al., 2016; Oda et al., 2005; Wiley and Burke, 2001). The EGFR gene is a major causative gene for several types of cancer; many drugs that target EGFR have been screened, identified, and developed (Ciardiello and Tortora, 2008; Murtuza et al., 2019). Some of these drugs show high therapeutic efficacy and are clinically approved to treat cancer (Murtuza et al., 2019; Sullivan and Planchard, 2016).

Target-based biochemical screening has a proven ability to identify molecular-targeted drugs, and indeed this approach has identified several EGFR-targeted candidates. However, target-based screening has produced only a limited number of drugs (Sams-Dodd, 2005). Many drug candidates failed in clinical trials due to either a lack of expected effects or the presence of unexpected effects. This attrition is due to, at least in part, inadequate knowledge of the target that is used for target-based screening (Hoelder et al., 2012). For example, if the target is required for cellular homeostasis, then inhibiting the target will result in adverse clinical results. The success of target-based screening is highly dependent on the initial selection of the target.

An alternative to target-based screening is phenotype-based drug screening, which has recently undergone a revival and re-evaluation (Johannessen et al., 2014; Moffat et al., 2014, 2017; Mullard, 2015; Wagner and Schreiber, 2016; Zheng et al., 2013). Although initial phenotype-based screening was associated with problems such as a lack of both quantitative evaluation and sufficient throughput, recent advances in automated microscopy and image analysis have solved these problems (Feng et al., 2009; Hughes et al., 2021; Simm et al., 2018; Smith et al., 2018; Young et al., 2008). Currently used phenotype-based screenings have several advantages over target-based screening. First, a compound identified in a phenotype-based screen exhibits a sought-for effect on the cellular process, and could be a good lead compound even if the mechanism of action (MOA) of the candidate is not known (Mullard, 2015). Second, since quantitative phenotype-based screening collects a huge amount of multi-parametric data in an unbiased manner, the use of adequate statistical approaches can identify novel MOA of known drugs (Feng et al., 2009; Tanabe, 2016). Third, phenotype-based screening could contribute to the repurposing or repositioning of existing drugs (Simm et al., 2018). Thus, using image-based profiling to screen pharmacologically active compound libraries is an efficient approach to evaluate the role of target molecules in the observed phenotype.

Previously, I developed an image-based compound profiling assay by evaluating the effects of compounds on EGFR-mediated signal transduction and EGFR membrane trafficking (Tanabe, 2016). In this assay, an EGFR inhibitor (CAS87912-07-8) and a microtubule polymerization inhibitor (nocodazole) produced a similar cellular phenotype, but other EGFR inhibitors did not produce this phenotype. A subsequent experiment showed that CAS87912-07-8 was a novel dual inhibitor of EGFR activity and microtubule polymerization. This result is one example of repositioning the MOA of a drug into a new category. Image-based phenotypic profiling can robustly predict the interaction between a compound and its target (Tanabe et al., 2018).

The aim of this study was to identify novel druggable targets associated with the EGFR signaling pathway using image-based profiling of a chemogenomic screening library. A lung-cancer cell line was treated with compounds from a chemogenomic screening library consisting of pharmacologically active compounds covering a broad range of cellular targets. Cells were then stimulated with EGF, and numerous features from images associated with activation of key signaling molecules in the EGFR pathway were extracted and used to classify the MOA of the compounds. Although many compounds have off-target effects that might be responsible for the observed phenotypes (Schenone et al., 2013; Whitebread et al., 2005; Xie et al., 2011), my system can infer the target responsible for the observed phenotypes induced by each compound. Using this system, I found that Rho-associated protein kinase (ROCK) and proteasomes are indispensable regulators of the EGFR pathway and are anticipated to be good candidate drug targets for cancers that involve EGFR signaling.

## Results

### 1. Visualization of signaling molecules to monitor the EGFR pathway

EGF activates various intracellular signals by coordinating intracellular membrane trafficking (Avraham and Yarden, 2011; Bakker et al., 2017). These signals can be visualized by immunofluorescence techniques. Fig. 1A shows representative responses after EGF stimulation of A549 cells (a lung-cancer cell line). At 5 min, EGFR exhibited a small punctate pattern in the cytoplasm, reflecting the localization of EGFR in transport vesicles. Between 30–60 min, EGFR signals appeared as larger early/late endosomes. At 180 min, the EGFR signals disappeared, indicating that EGFR had been completely degraded in lysosomes. The activation of two major signaling molecules in the EGFR pathway, Akt and ERK, were monitored by visualizing their phosphorylation status (pAkt and pERK, respectively) (Avraham and Yarden, 2011). A strong pAkt signal was specifically localized around the plasma membrane, whereas the pERK signal was observed throughout the cell (Fig. 1A). The intracellular distribution of EGF, which was visualized by fluorescence-conjugation, was similar to that of EGFR. This observation corresponds to previous findings that an EGF–EGFR complex is directed to lysosomes at high ligand concentrations (Sigismund et al., 2008). The intracellular distribution of transferrin, which cycles with its receptor between the plasma membrane and endosomes (Gruenberg et al., 1989; Mesaki et al., 2011), was different from that of EGF and EGFR. Internalized transferrin appeared as large vesicles at 5 min. Then at 30 min, small vesicles were observed in the perinuclear region, indicating that transferrin had moved to recycling endosomes. There was no signal detected at 180 min, indicating that transferrin was completely recycled back to the extracellular space. Some lipid kinases are activated by EGFR and are essential for producing proper cellular responses (Posor et al., 2015). However, the intracellular distribution of some products of these lipid kinases, namely PtdIns(3)P and PtdIns(4)P, did not significantly alter after EGF stimulation. This observation is most likely due to the presence of another pathway that produces these phospholipids (Roth, 1999, 2004). On the other hand, the amount of PtdIns(4,5)P2, a product of the lipid kinase PtdIns(4)P5K, significantly increased after the EGFR activation, consistent with the notion that PtdIns(4)P5K is activated by ligand stimulation of EGFR.

**Figure 1.**
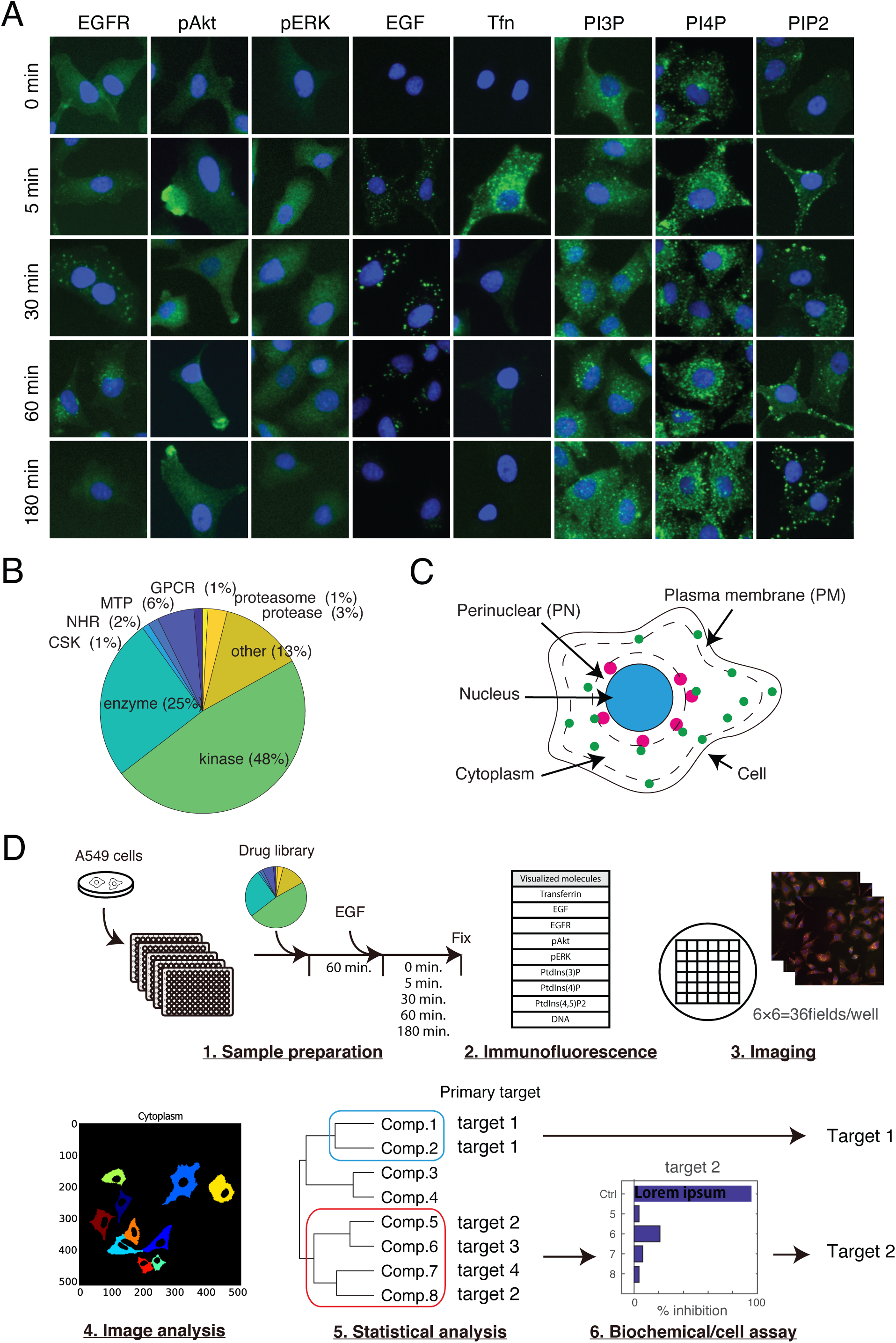
Overview of the experimental workflow for the image-based profiling. A. Visualization of signaling molecules to monitor the EGFR pathway. Representative images of molecules visualized in this study are shown. GFP-EGFR-expressing A549 cells were incubated with fluorescently labeled EGF or transferrin and processed for immunofluorescent procedures. Hoechst 33342 was used to identify the nucleus (blue). Each molecule is pseudo-colored green. B. The primary targets of the compounds in the library used in this study. C. Regions of interest (ROIs) were defined and used as the localization information for each signal. The signal of Hoechst 33342 and GFP-EGFR was used to define the nucleus and cell, respectively. D. An overview of the experimental workflow. Details are described in the Results.

### 2. Overview of the experimental workflow for the image-based profiling

The role of drug targets in the EGFR pathway was evaluated by quantitatively analyzing the molecules indicated in Fig. 1A at the indicated time points in the presence of compounds from a chemogenomic screening library consisting of 361 pharmacologically active compounds. The primary targets of the library are summarized in Fig. 1B and Supplementary Table S1. Briefly, the library consists of 172 kinase inhibitors, 11 protease inhibitors, three proteasome inhibitors, 92 inhibitors of other enzymes, four inhibitors of cytoskeleton formation, six nuclear hormone receptor inhibitors, 21 membrane transporter inhibitors, five GPCR inhibitors, and 47 inhibitors of other protein complexes. As most molecules visualized in this study dynamically change their intracellular localization after EGF stimulation (Fig. 1A), several subcellular regions of interest (ROIs) were defined and used as the localization information for each signal (Fig. 1C, see Experimental Procedures). All the image features acquired in this study are listed in Supplementary Table S2.

An overview of the experimental procedures is illustrated in Fig. 1D. Cells plated in 96-well plates were serum-starved for 5 h and then treated with compounds (at a final concentration of 10 µM) for 1 h. Cells were then stimulated with EGF (100 ng/ml), fixed after 0, 5, 30, 60, and 180 min, and processed for immunofluorescent staining. The cellular effects of nine molecules (the eight molecules described in Fig. 1A and Hoechst 33325) were visualized. Thirty-six fields per well were photographed (corresponding to an average of ∼300 cells) using an automated cell imager. The images were processed for image analysis to extract 134 image features (Supplementary Table S2). These image features were processed for principal component analysis (PCA) to reduce redundancies, and hierarchical cluster analysis was applied to compare the cellular effects of the compounds. Each cluster should contain several compounds that produce a similar phenotype. If all the compounds in a cluster share a single target, it implies that the target is responsible for producing the shared phenotype by that cluster of drugs (as represented by target 1 in Fig. 1D). If some compounds in a cluster do not share a common target (as shown in compounds 5–8), then two possibilities arise. One possibility is that all the targets (as represented by targets 2–4 in Fig. 1D) are indispensable for a specific pathway, and perturbations of these targets result in a shared phenotype. This could be confirmed in the literature and/or experimentally. A second possibility is that these compounds share a previously unrecognized specific target, which has been identified by the phenotypic screen. In the latter case, only one target might be responsible for producing the phenotype, and the role of this target could be confirmed by a biochemical assay (as represented by target 2 in Fig. 1D). Using the illustrated workflow, I evaluated the role of the compounds’ targets in the EGFR pathway.

### 3. The image-based profiling can detect characteristic phenotype from targeting a specific molecule or pathway

To quantitatively evaluate the phenotype induced by the compounds, 670 image features (134 features × 5 different time points) were obtained from cells treated with 361 compounds including DMSO as a control (the total number of image features obtained was therefore 670 × 361 = 241,870). The compounds’ effects were evaluated by a modified two-sample Kormogorov–Smirnov (KS) test, as previously described (Perlman et al., 2004; Tanabe, 2016). Acquired KS values were normalized using the standard deviations of control cells (Z-score, see Experimental Procedures), and Z-scores were used for subsequent analysis. Fig. 2A shows a heatmap of the Z-scores of 670 image features obtained from 361 compounds. Red and blue colors represent Z-scores that are higher and lower than Z-scores of control cells, respectively. As these image features share many redundancies (Fig. 2B, red and blue colors represent positive and negative correlation coefficients, respectively), PCA was used to reduce these redundancies, resulting in the identification of 361 principal components (PCs). For the first six PCs (ranked in order of weighting), the contributions of the representative image features to the EGF–EGFR pathway are shown as chart diagrams (Fig. 2C). PC1 contained image features that contributed highly to the EGF–EGFR pathway, and PC2 contained image features that contributed highly to pERK- and EGFR-associated signaling. On the other hand, PC3 and PC4 contained image features that contributed highly to pAkt- and PtdIns(3)P-associated signaling, respectively. These results indicate that each PC represents a distinct characteristic that contributes to the observed cellular phenotypes.

**Figure 2.**
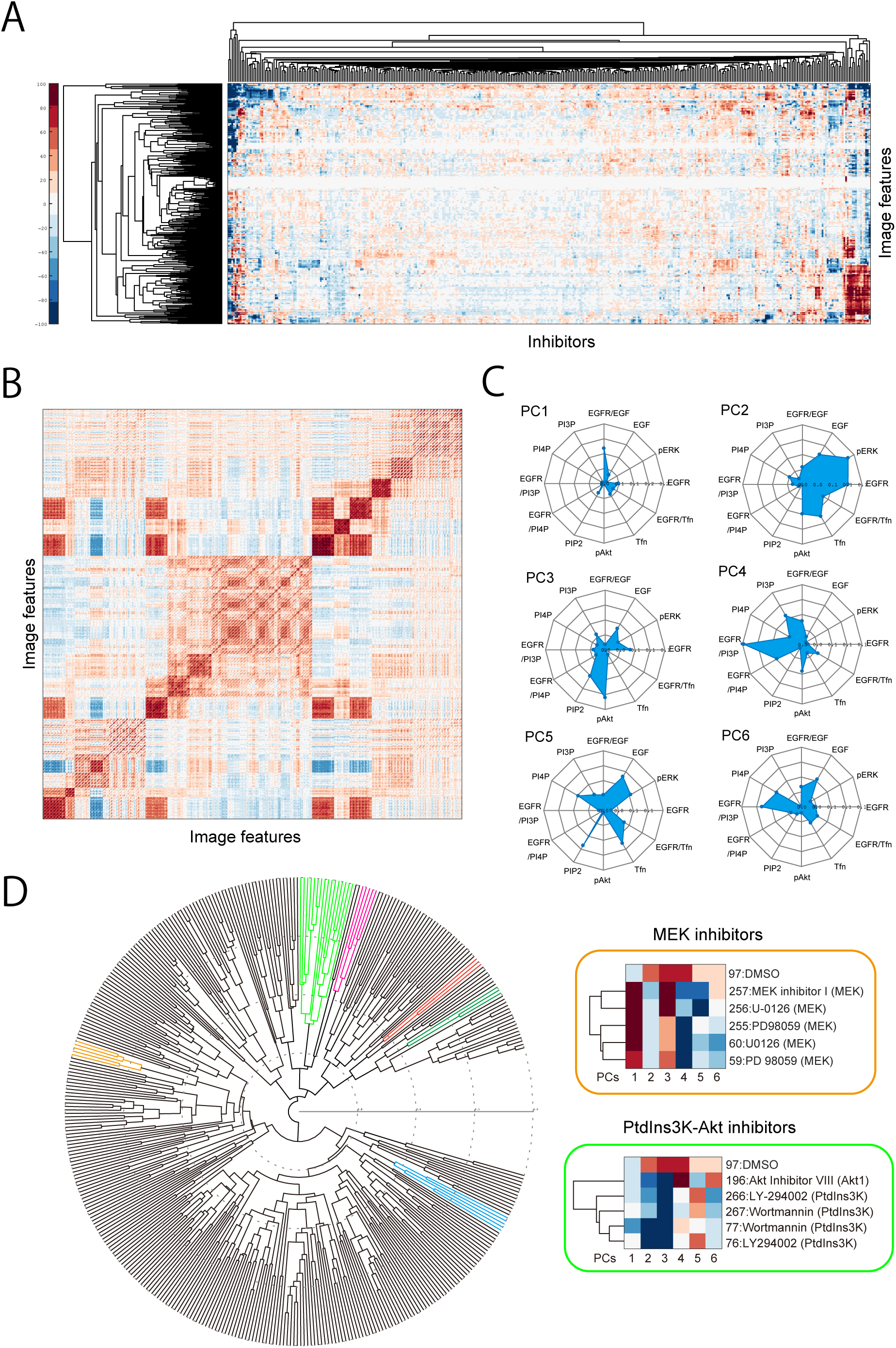
The image-based profiling can detect characteristic phenotype from targeting a specific molecule or pathway. A. A heatmap of 670 image features obtained from the phenotypic screening of 361 compounds. The image features listed in Supplementary Table S2 (134 image features) were collected at five time points and evaluated by a modified two-sample Kormogorov–Smirnov test. B. Self-correlation of image features shows that there are some strong correlations between image features. Red coloring indicates a positive correlation and blue coloring indicates a negative correlation. C. The contribution of representative image features in the first six principal components (PCs) are shown as char diagrams. D. Hierarchical cluster analysis of 361 compounds based on image features. The brown and a portion of the light-green cluster correspond to clusters containing well-known MEK inhibitors and PtdIns3K-Akt inhibitors, respectively. Clusters represented by other colors are described in Figs. 3–5.

Using all the PCs obtained, hierarchical cluster analysis was performed to evaluate target similarity and dissimilarity among the 361 compounds (Fig. 2D). As expected, several compounds known to share a common target formed a distinct cluster. For example, three MEK inhibitors (MEK inhibitor, U-0126 and PD98059) were found in a single cluster (light brown in Fig. 2D; several inhibitors were included in duplicate to confirm the accuracy of the experiment) (Alessi et al., 1995; Duncia et al., 1998; Wityak et al., 2004). Another example is that two PtdIns3K inhibitors (LY294022 and wortmannin) and one Akt inhibitor were found in another cluster (light green in Fig. 2D) (Arcaro and Wymann, 1993; Lindsley et al., 2005; Vlahos et al., 1994). This finding corresponds to the fact that both PtdIns3K and Akt have an essential role in the PtdIns3K–Akt signaling pathway (Oda et al., 2005), and reflects the cellular phenotype produced by each molecule during inhibition of the PtdIns3K–Akt pathway. Thus, the image-based profiling used in this study can detect characteristic phenotypes that result from targeting a specific molecule or pathway.

### 4. Repositioning of compounds that inhibit the PtdIns3K–Akt–mTOR pathway

Next, I focused on several clusters, which contain functionally unrelated compounds targeting distinct molecules. The cluster shown in Fig. 3A (corresponding to the light-green cluster in Fig. 2D) contained 14 compounds including four well-known inhibitors of EGFR signaling (LY 294002 and wortmannin, which inhibit PtdIns3K; Akt inhibitor VIII, which inhibits Akt; and torkinib (PP 242), which inhibits mTOR) (Arcaro and Wymann, 1993; Feldman et al., 2009; Lindsley et al., 2005; Vlahos et al., 1994). Thus, this cluster contained four compounds targeting the PtdIns3K–Akt–mTOR pathway (indicated in bold in Fig. 3A). The cluster also contained many compounds that, at least based on current knowledge, are not associated with effects on the PtdIns3K–Akt–mTOR pathway. For example, this cluster contained an Na channel inhibitor (amirolide), an inhibitor of geranylgeranyl transferase (GGTI-286), and a Cdk2/9 inhibitor (Benos, 1982; Lernert et al., 1995).

**Figure 3.**
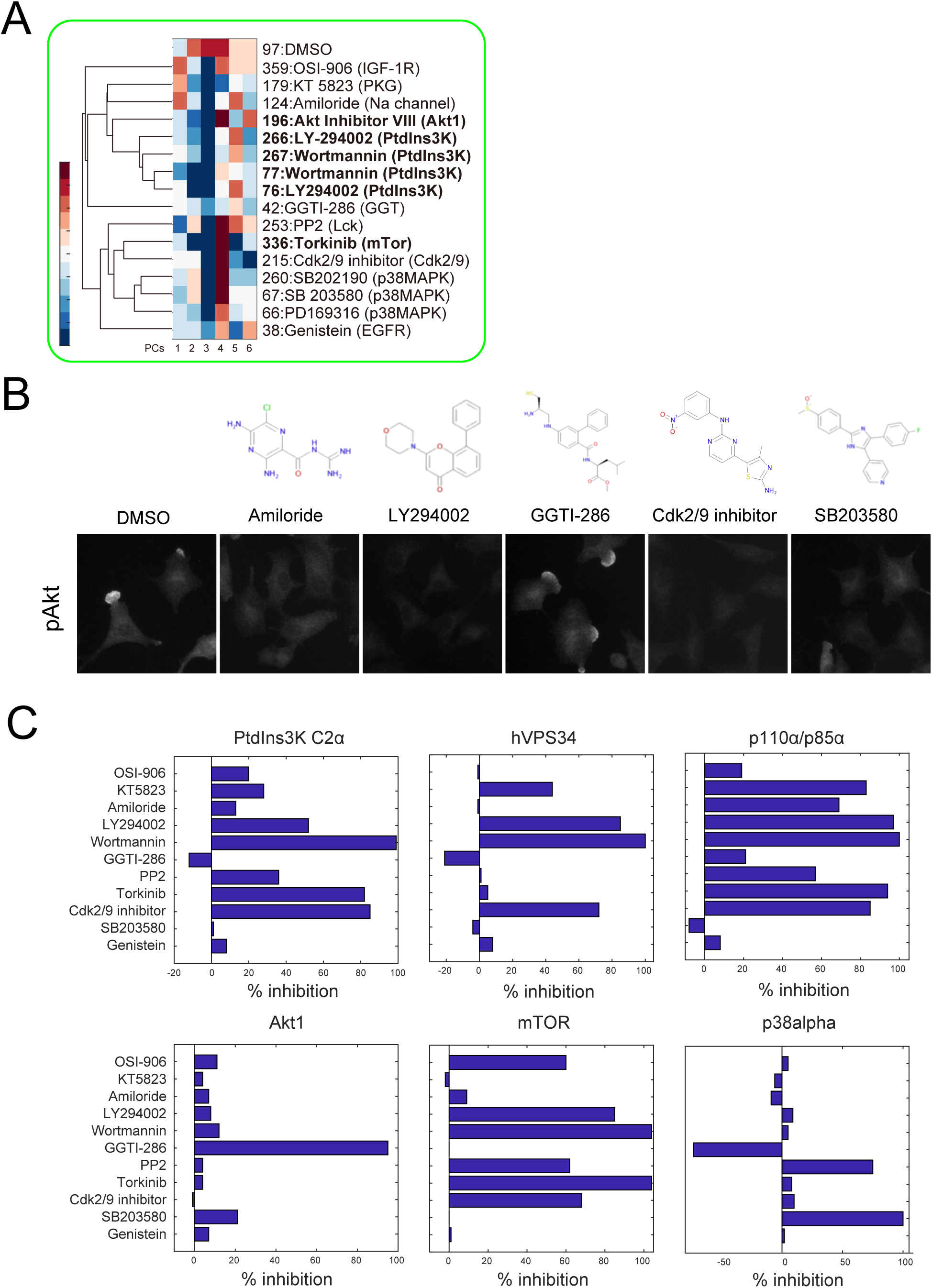
Repositioning of compounds that inhibit the PtdIns3K–Akt–mTOR pathway. A. A cluster containing PtdIns3K-Akt inhibitors, illustrated in light green in Fig. 2D, is shown with the contribution of the first six PCs. B. The chemical structure of several compounds and representative images showing the effect of these compounds on Akt phosphorylation. Note that these compounds have no structural similarity, but all of them except GGTI-286 inhibited Akt phosphorylation. C. An in vitro kinase assay showed that most compounds inhibited the PtdIns3K–Akt–mTOR pathway. The kinase activity of PtdIns3K class1-3, Akt, mTOR and p38a was measured in the presence of the listed compounds. Akt inhibitor VIII was omitted from this assay because this compound indirectly inhibits Akt by phosphorylation at Ser473. SB202190 and PD169316 were omitted from this assay as both compounds are analogs of SB203580.

When the cellular phenotypes represented by each PC (Fig. 3A) were carefully compared, the contribution of the third PC in images produced by all the compounds in this cluster was highly negative value opposite to positive in the control cells (note the relatively low change in cellular phenotype produced by GGTI-286 and genistein). This third PC contains image features that contribute highly to Akt signaling (Fig. 2C). Corresponding to this finding, the images (as shown in Fig. 3B, lower panels) show that the pAkt signal was remarkably suppressed in most of the compounds of this cluster (including both the known Akt inhibitor such as LY294002 and the unknown Akt inhibitors such as amiloride, Cdk2/9 inhibitor, and SB203580). On the other hand, GGTI-286 and genistein had limited effects on Akt phosphorylation (Fig. 3B and data not shown), reflecting the small effect observed on cellular phenotype in PC3 (Fig. 3A).

Although there is a possibility that targets other than ones from the PtdIns3K–Akt–mTOR pathway, such as the Na channel or geranylgeranyl transferase, have an indispensable role in the EGFR pathway, to the best of my knowledge there are no reports indicating such a possibility. Thus, it may be worth recalling that, in many cases, a compound can potentially affect multiple molecules in addition to its primary target (see Fig. 1D). To investigate if the compounds in the PtdIns3K–Akt pathway cluster possessed off-target activity, their activity was measured in an in vitro kinase activity assay (Fig. 3C). Surprisingly, almost all the compounds in the cluster showed significant inhibitory activity against Akt, PtdIns3Ks, or mTOR. For example, OSI-906 (an IGF-1 R inhibitor) inhibited mTOR; KT 5823 (a PKG inhibitor) and amiloride (an Na channel inhibitor) inhibited PtdIns3K class III (also known as hVPS34); and a Cdk2/9 inhibitor inhibited all classes of PtdIns3K (that is, PtdIns3K C2α, hVPS34, and p110α/p85α) and mTOR. Thus, most compounds in the cluster had inhibitory activity toward the PtdIns3K–Akt–mTOR pathway, and this inhibitory activity would contribute to the reduction in Akt phosphorylation observed in cells treated with these compounds (Fig. 3B). The geranylgeranyl transferase inhibitor GGTI-286 was found to be a direct inhibitor of Akt1 (Fig. 3C). This finding may explain why GGTI-286 did not reduce the Akt phosphorylation (Fig. 3B), yet produced a similar phenotype to PtdIns3K–Akt–mTOR pathway inhibitors that inhibit Akt activity by blocking Akt phosphorylation.

One compound in the cluster, SB203580 (a p38 MAPK inhibitor), inhibited neither PtdIns3Ks, Akt nor mTOR (Fig. 3C). Although three pyridinyl imidazole p38 MAPK inhibitors (SB203580, SB202190, and PD169316) were in this cluster, the structurally unrelated p38 MAPK inhibitor SB239063 was not included (Underwood et al., 2000). This finding suggests that these pyridinyl imidazoles reduce Akt phosphorylation through an unidentified target that is not the primary target of these compounds.

The above results show that the image-based profiling system could infer the non-primary target of the compounds, which might be responsible for the observed phenotype.

### 5. Identification of several essential roles of unvisualized targets – microtubules and actin

Next, I focused on two clusters, each of which appeared to share several unvisualized targets, that was not visualized using immunofluorescent technique. The first cluster contained nocodazole and vinblastine, both of which are well-known microtubule inhibitors (Fig. 4A, corresponding to the magenta cluster in Fig. 2D) (Jordan et al., 1992). This cluster also contained SB 225002 (a CXCR2 inhibitor), rotenone (an inhibitor of the mitochondrial electron transport), and PDGFR inhibitor IV (a PDGFR inhibitor), which do not share significant structural similarities (Fig. 4B, upper panels) (Chance and Hollunger, 1963; Ho et al., 2005; White et al., 1998). Fig. 4A shows there was a common perturbation in the fifth PC, which contains image features that highly contribute to transferrin-related activities (Fig. 2C). In accordance with this observation, treating cells with all the compounds inhibited the perinuclear localization of transferrin (Fig. 4B, middle panels). Inhibition of the perinuclear localization of transferrin is a characteristic feature of inhibition of microtubule polymerization, since it disrupts the intracellular transport of endocytic vesicles (Tanabe, 2016). Thus, microtubules were visualized in cells treated with these compounds, and all compounds tested significantly disrupted microtubule structure (Fig. 4B, lower panels). Based on the literature, SB 225002 and rotenone are known to inhibit microtubule polymerization in vitro (Brinkley et al., 1974; Goda et al., 2013; Meisner and Sorensen, 1966). Although further experiments are needed to show that PDGFR inhibitor IV is a novel inhibitor of microtubule polymerization, all compounds in this cluster have inhibitory activity against microtubules; this effect was not originally visualized using immunofluorescence techniques in the early part of this study (Fig. 1A).

**Figure 4.**
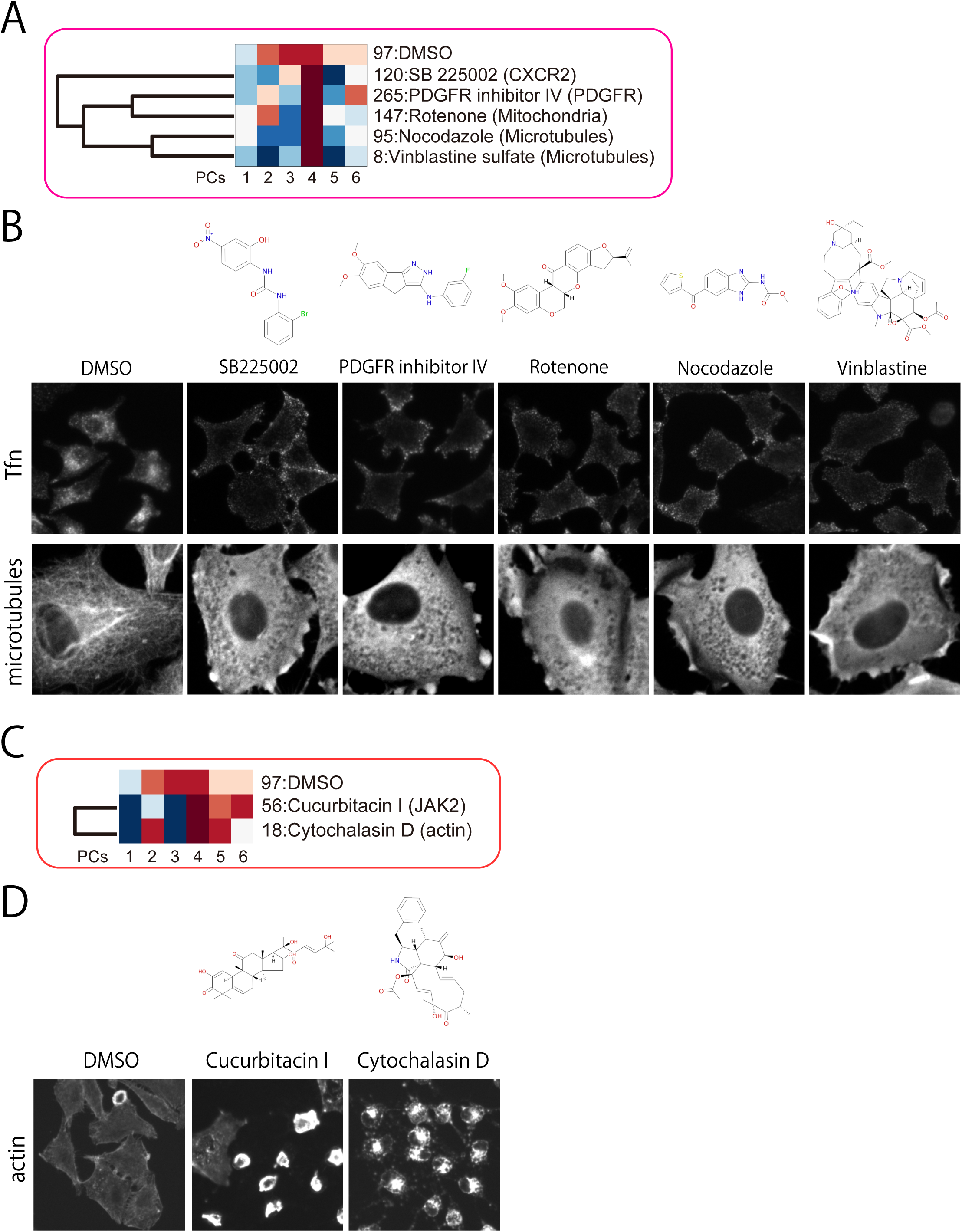
Identification of several essential roles of unvisualized targets – microtubules and actin. A. A cluster containing microtubule inhibitors identified from results presented in Fig. 2D (magenta). The contribution of the first six PCs is shown. B. Upper panels show the molecular structure of each compound. The middle panels show Alexa647-labeled transferrin, which was internalized following 30 min treatment with the compounds. The lower panels show microtubules visualized by anti-a-tubulin. C. A cluster containing an actin inhibitor identified from results presented in Fig. 2D (red) and shown with the first six PCs. D. The upper panels show the molecular structure of the compounds, and lower panels show the actin structure (visualized by phalloidin) following treatment with the indicated compounds.

Next, I focused on a cluster containing cytochalasin D, a widely used actin inhibitor (Schliwa, 1982), and cucurbitacin (a JAK2 kinase inhibitor; Fig. 4C, corresponding to the dark green cluster in Fig. 2D). These compounds have low structural similarity (Fig. 4D, upper panels) (Lee et al., 2010). Interestingly, cucurbitacin is also reported to inhibit actin (Knecht et al., 2010), and, as expected, actin structures were disrupted in cells treated with either compound (Fig. 4D, lower panels). Thus, the cluster containing cytochalasin D represents compounds characterized by the ability to depolymerize actin.

By investigating targets not visualized in Fig. 1A, I identified two clusters. Each cluster shares a common target: either microtubules or actin. Although investigating one of these clusters implied a new MOA of PDGFR inhibitor IV, the most important point indicated by the data presented in Fig. 4 is that microtubules and actin were both implicated in EGFR signaling because of the existence of clusters containing either microtubule or actin inhibitors. This finding is in agreement with the fact that microtubules and actin play essential roles in EGFR transport and signal transduction (Mesaki et al., 2011; Tanabe et al., 2011).

### 6. Identification of novel druggable regulators of the EGFR pathway

Finally, I focused on the two further clusters to identify novel indispensable regulators of the EGFR pathway. One cluster contained three ROCK inhibitors and the PKA inhibitor H-89 (Fig. 5A, corresponding to the blue cluster in Fig. 2D) (Chijiwa et al., 1990; Tamura et al., 2005; Uehata et al., 1997). H-89 is known to inhibit ROCK activity in vitro (Lochner and Moolman, 2006), so compounds in this cluster will produce their associated cellular phenotype when ROCK is inhibited. ROCKs are serine/threonine kinases activated by the small GTPase Rho that phosphorylate several substrates including LIMK, PTEN, and MLC (Amano et al., 2010). Although the association between Rho and EGFR is well known, the precise role of ROCK in the EGFR pathway has not been elucidated. Fig. 5A shows there was a common perturbation in PC1, which contains image features that highly contribute to EGF–EGFR signaling (Fig. 2C). So, I re-examined the EGF–EGFR signals produced by compounds in this cluster in ROCK-inhibitor treated cells. Fig. 5B shows the EGF signals 60 min after EGF stimulation. In DMSO-treated cells, EGF signals disappeared due to EGF/EGFR degradation in lysosomes, whereas a significant EGF signal was observed in ROCK inhibitor-treated cells. The phenotype observed during ROCK-inhibitor treatment implies that ROCK plays an essential role in EGF/EGFR degradation. ROCK has two isoforms, ROCK1 and ROCK2; the isoform responsible for EGF/EGFR degradation was examined by siRNA experiments. As shown in Fig. 5C, siRNA targeting ROCK1 inhibited EGF degradation as in the case of ROCK inhibitor-treated cells, but siRNA targeting ROCK2 did not produce this effect (Fig. 5C). A similar phenotype was also observed when siRNA targeting PTEN, a ROCK substrate, was used (Fig. 5C) (Li et al., 2005). These results indicate that the Rho–ROCK1–PTEN pathway regulates EGF/EGFR degradation.

**Figure 5.**
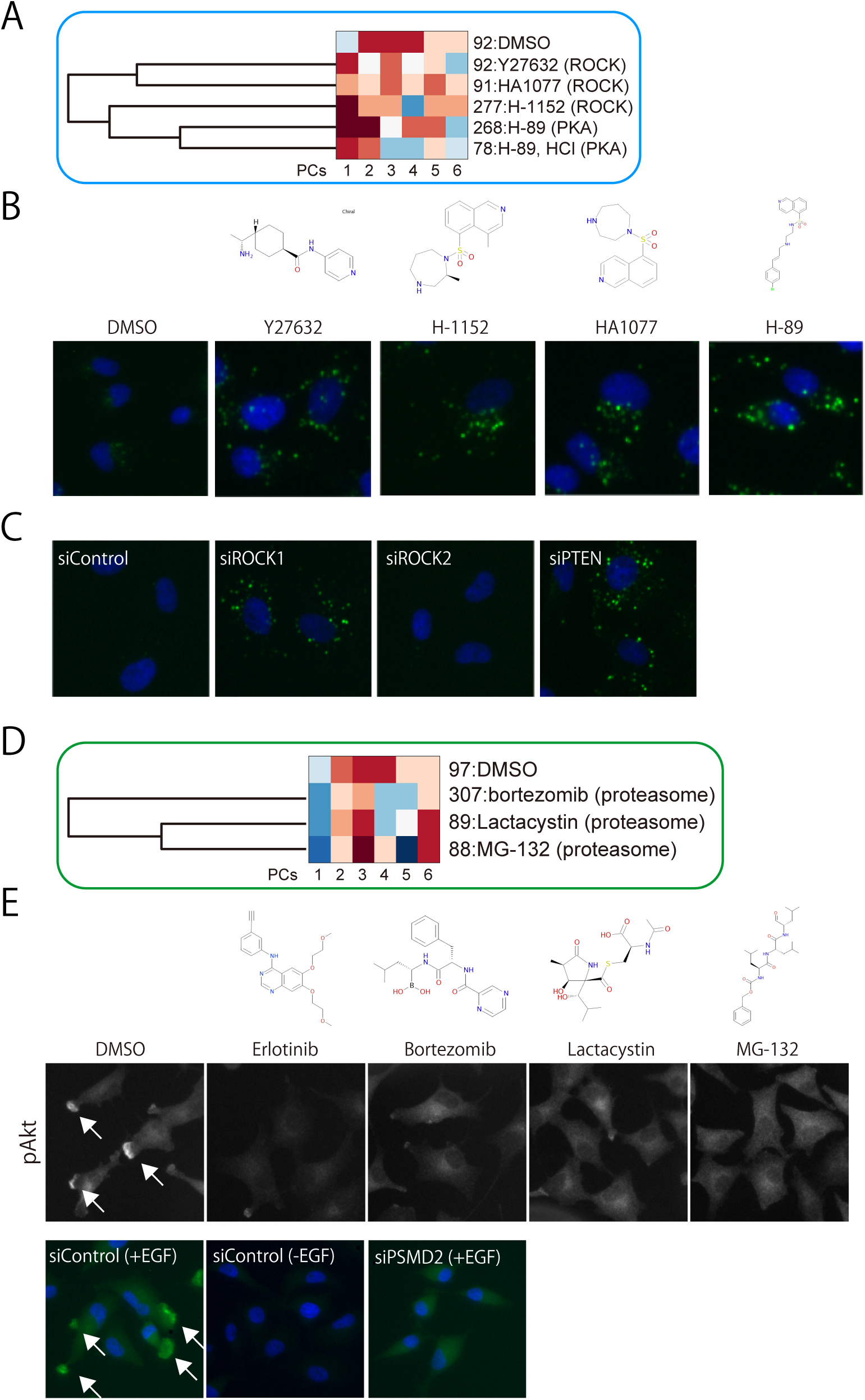
Identification of novel druggable regulators of the EGFR pathway. A. A cluster containing three ROCK inhibitors identified from results presented in Fig. 2D (blue) is shown with the first six PCs. B. The upper panels show the molecular structure of the compounds, and the lower panels show representative images of compound-treated cells. Images show Alexa647-labeled-EGF at 60 min after internalization (green) and Hoechst 33325 (blue). C. A549 cells were transfected with the indicated siRNAs. After 72 h transfection, cells were stimulated with Alexa647-labeled EGF (green), fixed after 60 min, and stained using Hoechst 33325 (blue). D. A cluster containing proteasome inhibitors identified from results presented in Fig. 2D (dark green) is shown with the first six PCs. E. The molecular structures of proteasome inhibitors are shown in upper panels. EGF was added to compound-treated cells for 30 min, and cells were fixed and processed for immunofluorescent procedures to visualize pAkt (middle panels). The arrows indicate pAkt signals at ruffle-like structures. Erlotinib, an EGFR inhibitor, was included to show the absence of pAkt signals. Note that cytosolic pAkt signals can be seen even in the presence of proteasome inhibitors. A549 cells were transfected with negative control siRNA (siControl) or PSMD2 siRNA (siPSMD2; lower panels). After 72 h, EGF was added to cells for 60 min (except for ‘siControl (-EGF)’), and then cells were processed for immunofluorescence assays using an anti-pAkt antibody (green) and Hoechst 33325 (blue).

Next, I focused on a cluster containing three proteasome inhibitors (Fig. 5D, corresponding to the red cluster in Fig. 2D) (Thibaudeau and Smith, 2019). In general, proteasomes degrade poly-ubiquitinated proteins independently of the endosome/lysosome pathway (Nandi et al., 2006; Piper and Lehner, 2011; Strous and Govers, 1999). Therefore, the possible involvement of proteasomes in EGFR signaling was surprising. A common perturbation in proteasome inhibitor-treated cells was observed in PC4 (Fig. 5D). PC4 contains image features that highly contributed to EGFR–PtdIns(3)P-, EGFR–PtdIns(4)P-, PtdIns(3)P-, and pAkt-related signaling. I compared fluorescent images of proteasome inhibitor-treated cells with control cells. Proteasome inhibitor treatment abolished the localization of pAkt signals in the sub-plasma membrane and the ruffle-like structure (peripheral pAkt; middle panels in Fig. 5E, arrows). However, cytoplasmic phosphorylation of Akt was retained in inhibitor-treated cells (in contrast to cells treated with the EGFR inhibitor erlotinib, in which Akt phosphorylation was not induced). These results, showing that that proteasome inhibitors inhibit peripheral pAkt, suggest that proteasomes play an essential role in the EGFR signaling pathway.

The 26S proteasome consists of a 20S catalytic core particle capped at both ends by 19S regulatory particles (Walters et al., 2004). Since three proteasome inhibitors (MG-132, bortezomib, and lactacystin) inhibit proteasome through different mechanisms (Thibaudeau and Smith, 2019), it is unlikely that inhibition of pAkt is due to a common off-target effect (e.g., MG-132 inhibits both calpains and cathepsin as non-primary targets) (Adams et al., 1998). To verify the significance of the proteasome in the phosphorylation of peripheral Akt, siRNA-based screening of 44 components of the proteasome was performed (see Supplementary Table S3 for the list of proteasome components). Of these proteasome components, PSMD2 siRNA significantly inhibited peripheral pAkt (Fig. 5E, bottom panels; see also Supplementary Figure S1). PSMD2 is a component of the 19S particle, and the result of this study corresponds to a previous immunoblot-based study showing that reduced expression of PSMD2 decreases cellular Akt phosphorylation (Matsuyama et al., 2011). PSMD2 depletion has a selective effect on some of the proteasome substrate (Li et al., 2018); thus PSMD2 might have a key role in the specific regulation of peripheral Akt phosphorylation.

## Discussion

In this study, numerous image features that reflect cellular phenotypes induced by compounds in a screening library were extracted and processed for statistical analysis to evaluate the role of the compounds’ targets in the EGFR signaling pathway. Drug discovery approaches that are based on a collection of small-molecule pharmacological agents and annotated targets, also called chemogenomic screening, require careful interpretation because most compounds or inhibitors have off-target activity in vivo. So, this study evaluated the effects of a compound library on quantitated cellular phenotypes, rather than on the known targets of the compounds. All compounds were evaluated and classified solely on phenotypic similarities. If several compounds share a common target, then these compounds are classified into the same cluster (Fig. 1D).

I first investigated whether the image-based system could identify compounds whose targets are directly related to molecules visualized by immunofluorescence techniques. As expected, three MEK inhibitors (MEK inhibitor I, U-0126, and PD98059) were classified into a single cluster, reflecting the notion that inhibition of MEK altered the phosphorylation status of ERK, which is a downstream substrate of MEK (Fig. 2D). Similarly, various PtdIns3K–Akt pathway inhibitors (Wortmannin, LY 294002, and Akt inhibitor VIII) were grouped in a cluster; compounds in this cluster all inhibit Akt phosphorylation. The above results indicate that the image-based system could evaluate cellular phenotypes induced by the inhibition of specific molecules. However, some clusters, containing both compounds known to target the EGFR pathway and other compounds, are thought to act independently of the EGFR pathway. For example, the cluster shown in Fig. 3A included several inhibitors targeting PtdIns3K, Akt, or mTOR, yet also contained inhibitors targeting IGF-1R, an Na channel, geranylgeranyl transferase, and other molecules. However, a biochemical assay revealed that such seemingly unrelated compounds also had inhibitory activity against PtdIns3K, Akt, or mTOR. Thus, the compounds in this cluster produce common cellular phenotypes by inhibiting the activity of the PtdIns3K–Akt–mTOR pathway.

Further, this image-based approach is applicable to molecules not visualized by immunofluorescence techniques. Clusters shown in Fig. 4A and C contain inhibitors of microtubules and actin, respectively. The cytoskeleton was not visualized in this study, but the presence of a cluster of cytoskeleton inhibitors infers that the cytoskeleton is important in EGFR and/or transferrin transport. This is a reasonable assumption, as microtubules and actin are both essential for the transport of EGFR and transferrin (Mesaki et al., 2011; Tanabe et al., 2011). This result suggests that the image-based system can identify targets that have an essential role in the EGFR pathway, even if the target itself is not directly measured.

In this study, I focused on two clusters sharing a target whose role on the EGFR pathway was not yet established. One cluster contained four inhibitors of ROCK, which is a well-known Rho effector molecule (Amano et al., 2010). I found that the degradation of EGF/EGFR was significantly inhibited in cells treated with ROCK inhibitors (Fig. 5B). A similar phenotype was observed in cells treated with ROCK1 siRNA but not in cells treated with ROCK2 siRNA (Fig. 5C). ROCK1 and ROCK2 are both highly expressed in A549 cells (Vigil et al., 2012) and share many substrates in vitro, but each isoform might have a unique role in vivo (Loirand, 2015). Interestingly, the expression of ROCK1 is associated with a poor prognosis in various types of cancer, including NSCLC (Akagi et al., 2014; Bottino et al., 2014; Hu et al., 2019; Liu et al., 2011; Tang et al., 2019); indeed, several ROCK inhibitors are used clinically (Feng et al., 2016; Olson, 2008; Rath and Olson, 2012). Further, reduced expression of PTEN, a ROCK substrate, also inhibited EGFR degradation (Fig. 5C). These results suggest that ROCK1 and PTEN have an indispensable role in mediating EGFR degradation. Rab7, a well-known essential regulator of the endosome–lysosome system, is a substrate of PTEN (Shinde and Maddika, 2016). Thus, a novel notion emerges that the Rho–ROCK1–PTEN–Rab7 axis promotes EGFR degradation.

Another cluster (Fig. 5D) consisted of three proteasome inhibitors (bortezomib, lactacystin, and MG-132). To the best of my knowledge, the involvement of the proteasome in EGFR signaling has not been reported. In this study, proteasome inhibitors significantly suppressed periphery-localized Akt phosphorylation (indicated as arrows in DMSO-treated control cells; Fig. 5E). The proteasome is generally involved in degrading ubiquitinated proteins, although there are some exceptions (Erales and Coffino, 2014). There is a report indicating that K63-linked ubiquitination regulates Akt activation (Yang et al., 2013). K63-linked ubiquitination acts as a switch in signal transduction rather than as a signal for degradation by the proteasome, but there are some reports that ubiquitinated pAkt also recruits the proteasome complex (Fan et al., 2013). In this study, I conducted experiments using a siRNA library against proteasome-related genes and identified PSMD2, whose reduced expression decreased peripheral Akt phosphorylation as proteasome inhibitors. This finding is in agreement with a previous study showing that PSMD2 is required for cellular Akt phosphorylation by balancing activation of p38MAPK and Akt (Matsuyama et al., 2011). Several reports indicate that PSMD2 expression correlates with poor prognosis in breast cancer and lung adenocarcinomas (Li et al., 2018; Matsuyama et al., 2011). On the other hand, several proteasome inhibitors are in clinical use, but their precise MOA remains to be elucidated. PSMD2, the non-catalytic proteasome subunit, might be a good target for anticancer therapy.

The main purpose of this study was to identify novel druggable regulators of the EGFR pathway using an image-based phenotypic assay to screen a pharmacologically active compound library. Although the use of chemical library assays is associated with potential problems, such as the off-target activity of the compounds, the system used in this study can infer the target responsible for a cellular phenotype by combining the results of unsupervised machine learning and a biochemical assay. The regulators of the EGFR pathway identified here could be druggable targets, and lead compounds might be identified following drug development and repurposing. The image-based approach used here could be easily applied to other signal pathways and cellular responses, and would be expected to find essential druggable regulators of each pathway. Further, comparing the chemical profile of each compound obtained under different experimental conditions could be used to identify a specific regulator in each signaling pathway.

## Experimental Procedures

### Reagents and antibodies

Human recombinant EGF (236-EG) and human transferrin (2474-TR) were purchased from R&D Systems. Alexa Fluor 647-conjugated EGF (E35351), Alexa Fluor 647-conjugated human transferrin (T23366), and Hoechst 33342 (H3570) were purchased from Invitrogen. Rabbit anti-phospho-ERK (Thr202/Tyr204) monoclonal antibody (#4370) and rabbit anti-phospho-Akt (Ser473) monoclonal antibody (#4060) were purchased from Cell Signaling Technologies. Mouse anti-PtdIns(4)P monoclonal antibody (Z-P004) and mouse anti-PtdIns(4,5)P2 monoclonal antibody (Z-P045) were purchased from Echelon. Rabbit anti-GST polyclonal antibody (SC-459) was purchased from Santa Cruz Biotechnology. Mouse anti-tubulin antibody (T5168) was purchased from Sigma. All secondary antibodies were purchased from Invitrogen. All antibodies were preserved at −30 °C after the addition of an equal volume of glycerol. The pharmacologically active compound library (SCADS1-4) was obtained from the Screening Committee of Anticancer Drugs (Japan). Recombinant GST-HrsFYVE protein was prepared as described previously (Henmi et al., 2016).

### Compound treatment and EGF stimulation

A549 cells expressing GFP-tagged EGFR were obtained from Sigma. Cells were plated on Edge plates (Thermo Scientific) in DMEM containing 10% serum and antibiotic–antimycotic solution (Sigma), and incubated at 37°C, 5% CO_2_ for at least 16 h. Then, cells were serum-starved for 6 h by replacing the medium with DMEM containing 0.1% BSA, and were treated with compounds at 10 µM for 1 h prior to ligand stimulation. Cells were stimulated with fluorescently labeled or unlabeled ligands (see Supplementary Table S4 for the ligands used). The fluorescently labeled ligands were replaced by unlabeled ligand 5 min after the initial ligand stimulation. At the indicated times after the stimulation (0, 5, 30, 60, and 180 min), cells were fixed by the addition of an equal volume of 4% paraformaldehyde (09154-85, Nacalai Tesque) for 15 min, followed by washing in PBS three times. The fixed cells were subsequently processed for immunostaining.

### Immunofluorescence

The immunofluorescence procedures were performed as described previously (Tanabe, 2016). Briefly, cells were permeabilized with ice-cold methanol containing either 0.3% Triton-X100 or digitonin according to the primary antibodies used, and then blocked with 0.1% bovine serum albumin. GST-HrsFYVE protein was added to the blocking buffer for cells that were to be labeled with PtdIns(3)P antibodies, incubated for 1 h at room temperature, and washed with PBS three times. Cells were incubated with the following primary antibodies for 1 h at room temperature: anti-pERK (1:1000), anti-pAkt (1:2000), anti-PtdIns(4)P (1:2000), anti-PtdIns(4,5)P2 (1:1000), or anti-GST (1:200). Cells were washed with PBS twice and subsequently incubated for 1 h at room temperature with the appropriate secondary antibodies (1:250) and Hoechst 33432. Cells were washed twice with PBS, fixed with 2% paraformaldehyde for 5 min at room temperature, washed with PBS, and stored at 4 °C until cell images were photographed.

### Image acquisition and analysis

Stained cells were photographed using a cell image analyzer (CellInsight, Thermo Scientific) equipped with a 20× objective lens. Hoechst and GFP-EGFR signals were used in all marker sets to define the ‘nucleus’ and ‘cell’, respectively. In each well, 36 fields were photographed. Image analysis was performed using the CellProfiler platform (Broad Institute) (Carpenter et al., 2006). ‘Cytoplasm’ was defined by subtracting ‘nucleus’ from ‘cell’. The ‘nucleus’ region was expanded by 5 pixels, and the resultant region was named the ‘perinuclear’ region. The ‘cell’ region was shrunk by 7 pixels, and the resultant region was named the ‘plasma membrane’ region. The signal intensities and the number of signals per cell were measured from each ROI. Image features measured in this study are listed in Supplementary Table S2.

### Statistical analysis

Data were stored in a local MySQL server and processed for statistical analysis using MatLab (Mathworks). The selected image features (known as descriptors) of each compound were compared with those observed in control (DMSO-treated) cells using the one-sided, signed two-sample KS test as previously described (Perlman et al., 2004; Tanabe, 2016). Signed KS statistics were standardized using control standard deviations (referred to as a Z-score). All PCs were used for subsequent hierarchical clustering using the correlation distance and the average linkage.

### Biochemical assays

Kinase activity assays were performed using the SelectScreen biochemical kinase profiling service provided by Life Technologies. Adapta screening protocols were used to measure PIK3C2A (PtdIns3K-C2α), PIK3C3 (hVPS34), and PIK3CA/PIK3R1 (p110α/p85α) activity. Z’-LYTE screening protocols were used to measure AKT1, FRAP1 (mTOR), and MAPK14 (p38α, direct activity) activity. All compounds were assayed at 10 µM. GGTI-286 was used at 0.5 nM–10 µM in an assay to measure concentration-dependent activity. The percentage inhibition produced by each compound was calculated as the mean value of two independent measurements.

### siRNA transfection

Cells were transfected with specific siRNAs using Lipofectamine RNAiMAX (Life Technologies) using reverse transfection according to the manufacturer’s instructions. siRNAs against ROCK1, ROCK2, and PTEN were purchased from Sigma. A custom-made proteasome siRNA library was purchased from Bioneer. At 72 h after transfection, cells were either incubated with Alexa488-labeled EGF or processed for immunofluorescent procedures.

## Acknowledgments

The author thanks M. Satake for reviewing the manuscript, Y. Henmi for his technical assistance and members of my institute for their kind supports. The SCADS inhibitor kit was provided by the Screening Committee of Anticancer Drugs supported by a grant-in-aid for Scientific Research on Innovative Areas and Scientific Support Programs for Cancer Research from the Ministry of Education, Culture, Sports, Science and Technology, Japan. This work was supported by grants from the Ministry of Education, Science, Sports, Culture and Technology of Japan (23770148, 23113721, and 25670149) and the Takeda Science Foundation.

## Author Contribution

K.T. conducted all the experiments and wrote the paper.

## Declaration of Interests

The author declares no competing interests.

## Notes

### Competing Interest Statement

The authors have declared no competing interest.

