## Supplementary Figure S1 for "Image-based phenotypic profiling of a chemogenomic screening library identifies novel druggable targets in the EGFR-pathway"

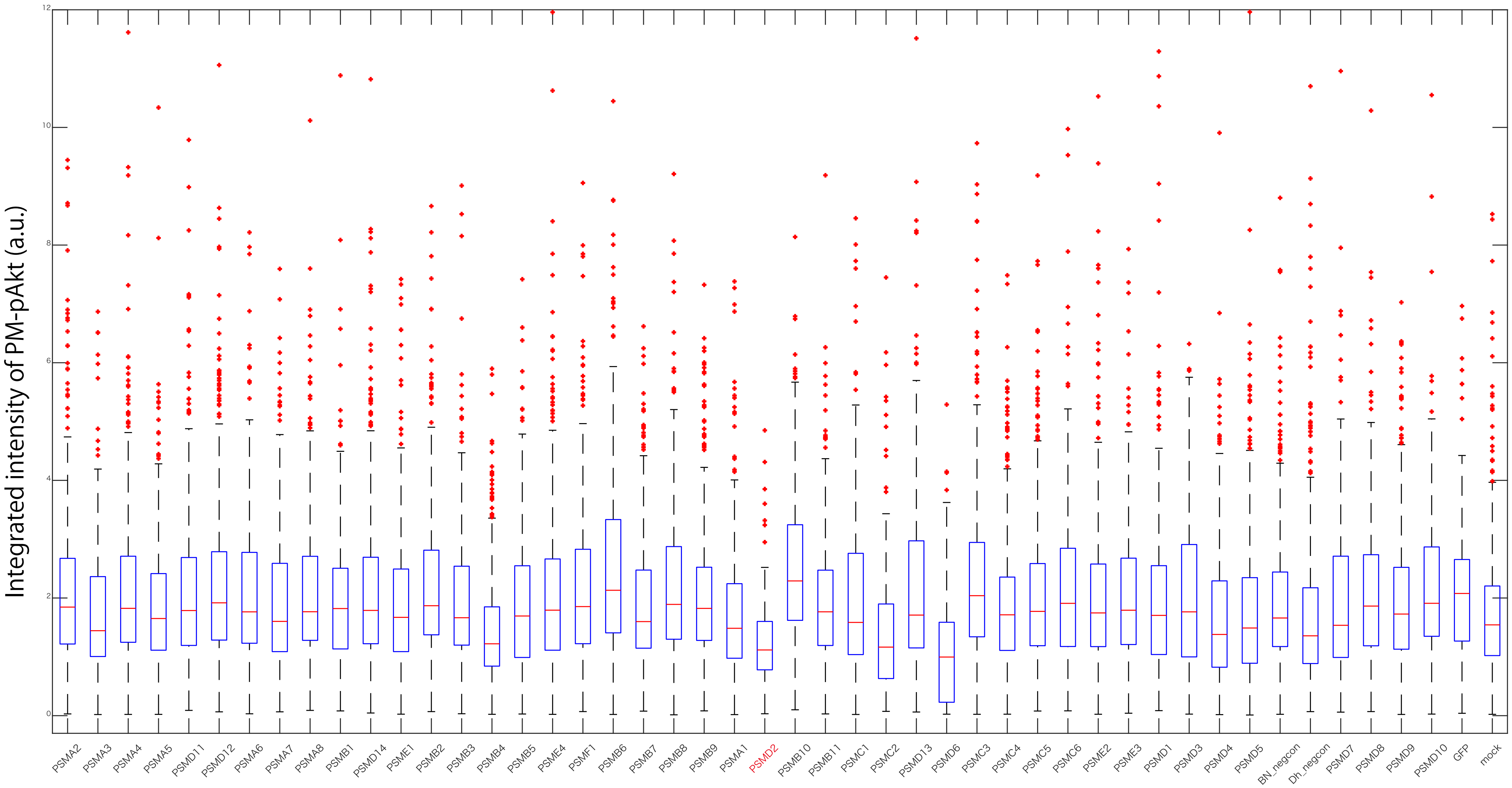


**Supplementary Figure S1. A proteasome siRNA library identifies PSMD2 as an indispensable gene for Akt phosphorylation.** Proteasome siRNAs were transfected into A549 cells expressing GFP-EGFR using Lipofectamine RNAiMAX. After 72 h transfection, Alexa488-labeled EGF was added to cells for 30 min. The cells were then fixed and stained with an anti-pAkt (Ser473) antibody and Hoechst 33432. Thirty-six images were taken from each sample and analyzed using CellProfiler software. The integrated intensity of pAkt signals around the sub-plasma membrane of each cell was quantified.
